# Stochastic Gene Expression Model with State-Dependent Protein Activation Delay

**DOI:** 10.64898/2026.03.31.715756

**Authors:** Poulami Chatterjee, Abhyudai Singh

**Affiliations:** Department of Electrical and Computer Engineering, University of Delaware, Newark, DE, USA; Departments of Electrical and Computer Engineering, Biomedical Engineering, Mathematical Sciences, and the Center for Bioinformatics and Computational Biology, University of Delaware, Newark, DE, USA

## Abstract

Cells must maintain stable protein levels despite the inherently stochastic nature of gene expression, as excessive fluctuations can disrupt cellular function and impair the reliability of decision-making. Regulatory mechanisms, such as negative feedback, buffer protein fluctuations. Yet, it remains unclear how fluctuations are affected by delays that depend on a molecule’s specific state. Here, we develop a stochastic model in which proteins are produced in bursts as inactive molecules and pass through a series of intermediate steps before becoming active. The duration of such activation delays depends on the current level of active protein, creating a state-dependent feedback loop. Our model provides explicit analytical expressions relating the delay structure and feedback strength to the variability of active protein levels, quantified using the Fano factor, and shows that state-dependent delays can reduce fluctuations below the baseline expected from simple bursty production. Stochastic simulations confirm these predictions, and incorporating negative feedback in burst production further decreases variability while keeping system behavior predictable. These results reveal how temporal and state-dependent regulation stabilizes protein expression, offering guidance for understanding natural cellular control and designing robust synthetic gene circuits.

## I. Introduction

Protein synthesis in a cell is a fundamental process that includes transcription and translation, collectively referred to as gene expression. In this process, genes are transcribed into mRNA, and the mRNA gets translated into protein, allowing the conversion of genetic information from genes into protein molecules. Protein levels can fluctuate within a homogenous population of genetically identical cells because protein production mechanisms are inherently unpredictable. Studies have shown that variability in protein production arises from both intrinsic and extrinsic sources. Intrinsic noise originates from random fluctuations in transcription and translation [1]–[7]. Extrinsic noise, on the other hand, stems from differences in cellular stage, cell growth, and the surrounding microenvironment [8]–[13]. Differences between cells arising from stochastic gene expression [14]–[18] driven by mechanisms such as transcriptional and translational bursting, variability in mRNA and protein decay rates, delayed protein production, and fluctuations in regulatory processes play an important role in biology and medicine. These differences directly affect cell fate [19]–[23], microbial bet hedging [24], [25], and drug resistance in bacteria and cancer [26]–[32]. Quantifying and modeling these fluctuations is therefore essential for understanding cellular behavior and for designing synthetic gene circuits with predictable dynamics.

Despite the natural randomness in gene expression, cells can maintain stable function and homeostasis. They achieve this by using various regulatory mechanisms that reduce fluctuations, or noise, in the level of a given protein. One common strategy is negative feedback, in which a protein inhibits its own synthesis, either directly or indirectly [33]–[35], [35]–[38], [38]–[42]. Recent theoretical work on synthetic control mechanisms, such as PID control schemes, has offered ways to fine-tune protein levels and further reduce variability [43], [44]. Another important mechanism for fluctuation reduction involves decoy binding, where transcription factors are sequestered by nonfunctional sites, buffering fluctuations in active protein levels [45]–[47]. Similarly, mRNA transport from the nucleus to the cytoplasm can smooth out bursts of gene expression, reducing variability in protein levels [48], [49]. Recent studies also show that active proteins can directly affect burst-like production, leading to state-dependent delays in expression [50]–[53]. This delay is due to the fact that the availability of active molecules, such as transcription factors and ribosomes, modulates the size of transcriptional and translational bursts.

In contrast to gene expression models without delay, an important open question is how state-dependent delays influence the stochastic dynamics of gene expression. Specifically, it is still unclear if delays increase or decrease variations in protein levels, particularly when the delay is dependent on the system’s molecular state. To address this question, we develop a stochastic model in which an inactive protein is converted into an active protein through an arbitrary number of delay stages, *n*. Biologically, these stages can represent intermediate processes such as transcription, translation, protein folding, or intracellular transport that occur before the final protein becomes active. State dependence is incorporated by allowing the transition rates between delay stages to depend on the active protein level.

In biochemical systems, state-dependent delays naturally occur when the timing of a reaction or transition relies on the system’s state, such as the quantity of an active protein or enzyme. In gene expression, such delays can occur when an inactive protein needs to undergo multiple activation steps or interact with regulatory molecules before becoming functionally active, such as through complex formation, transport mechanisms, or enzymatic modification. Our technique incorporates state dependency by representing transition rates across the delay stages as explicit functions of active protein abundance. This allows the effective delay to fluctuate dynamically with the molecular state instead of remaining constant.

We can capture these mechanistic characteristics and give a more accurate explanation of protein kinetics by including state-dependent delays into stochastic models. While effective simulation frameworks for chemical systems with time- and state-dependent propensities were developed in [55], earlier work introduced stochastic delay processes that depend on reaction history or system state [56]. Reviews of stochastic and delayed stochastic gene expression models highlight the importance of explicitly modeling transcriptional and translational delays in regulatory networks [57]. Oscillatory gene regulation has been studied using discrete stochastic delay models [58], [59], and analytical distributions for protein and mRNA levels in delayed gene expression systems—extendable to state-dependent formulations—have been derived in [60]. More recently, multistep biochemical reaction models have demonstrated how effective delays arise as functions of system state [50], and explicit state-dependent transcriptional and translational delays have been examined in gene regulatory networks [61]. Stochastic analysis of feedback control with delays further highlights the influence of such delays on network dynamics [62]. All of these results support the use of state-dependent delays in our stochastic model and set the groundwork for understanding how they affect fluctuations at the protein level.

We employ two complementary approaches to characterize fluctuations in the active protein dynamics. First, we employ the Stochastic Simulation Algorithm (SSA) [54] to construct accurate stochastic trajectories of the system across time, capturing the full impacts of inherent fluctuations. Second, we apply the Linear Noise Approximation (LNA) [63]–[66], which provides an analytical approximation to stochastic fluctuations around the steady state. Using these methods, we quantify how fluctuations in the active protein depend on the average delay at steady state. We aim to compare a no-delay system with a multi-delay system in order to assess the impact of delays on fluctuation characteristics. For comparison, we first review the commonly used no-delay protein production model, in which the protein is activated instantaneously. This schematic is shown in the top panel of Fig. 1(A). The one-step delay model is then used to determine fluctuation levels and validate our analytical results using SSA. Next, we proceed to include an additional control factor, negative feedback in burst rate, with the one-step state-dependent delay model. Our analysis shows that, despite these factors, the fundamental fluctuation limit remains unchanged and can even be reduced below that of a system without delays.

**Fig. 1:**
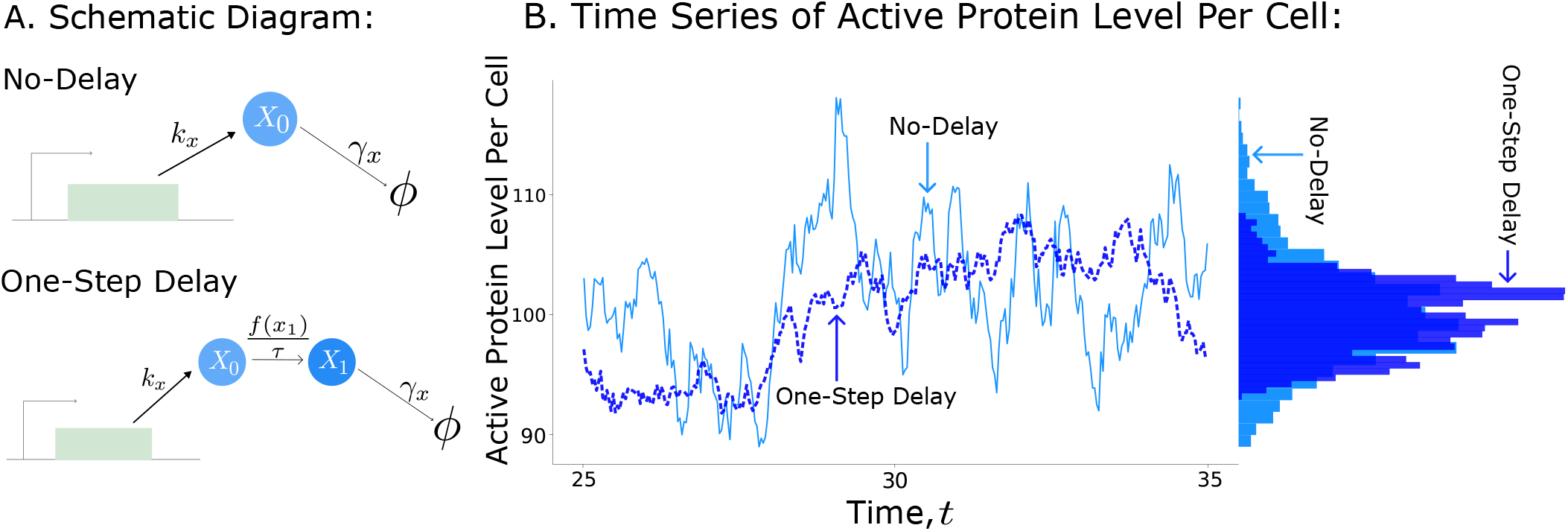
State-dependent protein activation delay decreases fluctuations in active protein levels. **A**. The left panel illustrates two protein production models: a no-delay system (top), in which proteins are produced in bursts and degrade immediately, and a one-step state-dependent delay system (bottom), where inactive proteins (*X*_0_) convert to active proteins (*X*_1_) after a state-dependent delay at a rate *f*(*x*_1_)*/τ* . *τ* is the average time delay and *f*(*x*_1_) is the state-dependent delay function of the form in Eq. (19). **B**. The right panel shows the time series of both systems generated using the Stochastic Simulation Algorithm [54]. The no-delay model is shown in light blue (solid), and the one-step delay model is in deep blue (dashed). The corresponding histograms of protein levels reveal reduced fluctuations in the one-step delay system compared to the no-delay system. For the results in Panel B, the burst size is *B* = 10, and the burst rate is *k*_*x*_ = 10 events per mean protein lifetime. For the SSA results in Panel B, the burst size is *B* = 10 and the burst rate is *k*_*x*_ = 10 per unit time, with a decay rate *γ*_*x*_ = 1 and an average delay *τ* = 2, which corresponds to approximately 2.88 protein half-lives.

The paper is organized as follows. In Section II, we introduce the fundamental model and compare active protein levels in the no-delay system with those in the one-step, state-dependent delay system II. We then investigate the one-step delay model in Section III, and extend the analysis to models with a number of delay stages, *n*, in Section IV and compare the results. Finally, we extend the analysis to models including feedback in burst frequency in a one-step delay model and with an arbitrary number of delay stages, *n*, in Section V.

## II. Model Formulation: No-Delay System

Transcriptional and translational processes can result in burst-like protein production, in which proteins are generated in short, unpredictable bursts rather than continuously. These bursts can be caused by either irregular mRNA transcription or bursts of translation from existing mRNA molecules [20], [67], [68]. In such systems, protein molecules are created in discrete stochastic events, resulting in high variability in protein abundance [14], [69]. Proteins are created in our model by burst-like synthesis events, following which they go through a delayed activation phase before fading. The delay indicates the intermediary stages required for the protein to become functionally active, whereas stochastic burst creation is a significant source of intrinsic noise in the system [15], [70].

To model these fluctuations, we start with the simplest stochastic system, which does not include any delay. In this system, the protein is activated instantaneously, and it is therefore referred to as the no-delay system. This model has been extensively studied in previous works [2], [34], [71] and serves as a baseline for evaluating the effects of temporal delays or regulatory feedback. We begin by reviewing this framework, in which proteins are produced in stochastic bursts and subsequently degraded.

The protein species is denoted by *X*_0_ and the level of protein at time *t* is denoted by *x*_0_(*t*). Protein production occurs in bursts according to a Poisson process with a burst rate or frequency *k*_*x*_. Each burst event generates *B* protein molecules, where *B* is an independent and identically distributed (i.i.d.) random variable with probability mass function

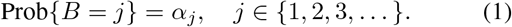

The probability of a burst event occurring in an infinitesimal interval (*t, t* + *dt*) that increases protein counts by exactly *j* molecules is *k*_*x*_*α*_*j*_*dt*. Each protein molecule degrades at a constant rate *γ*_*x*_, resulting in the probability of degradation event occurring in (*t, t* + *dt*) being *γ*_*x*_*x*_0_(*t*)*dt*. Table I provides an overview of the stochastic model and a schematic of the process is shown in Fig. 1(A). We examine the statistical moments of the protein distribution to quantitatively describe the following stochastic dynamics. Moments are statistical quantities used in probability and stochastic processes to characterize various key features of a random variable [72]–[74]. To study fluctuations, we begin by examining the differential equations that describe the time evolution of these statistical moments [75], [76]. Using Table I, the differential equations governing the expected value of any differentiable function, *ϕ*(*x*_0_), are given by:

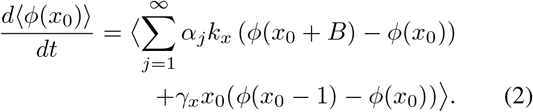

**TABLE I:**
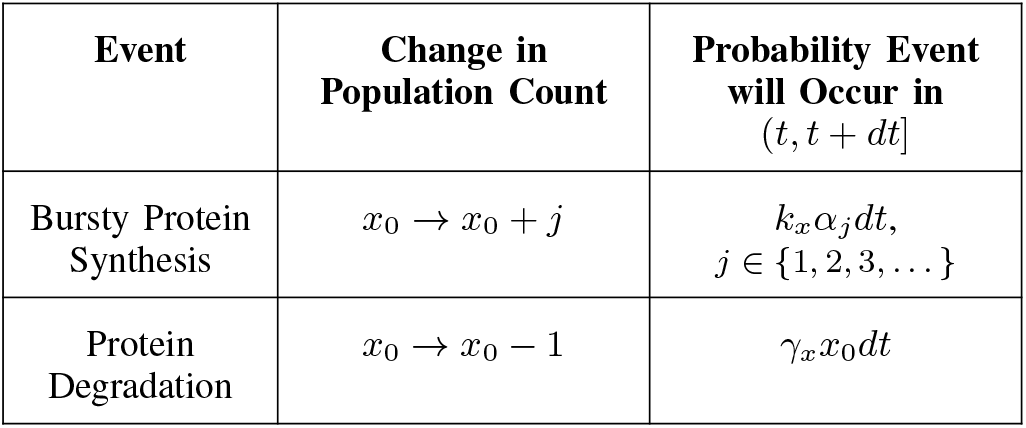
Stochastic model of bursty gene expression with instantaneous protein activation (no-delay).

Throughout this work, ⟨·⟩ denotes the expected value of the random process, while 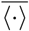 represents the corresponding steady-state expectation. The system’s moment dynamics are obtained by choosing

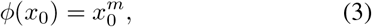

where *m* = 1, 2, 3 … gives the first, second, and higher-order moments, respectively, as follows:

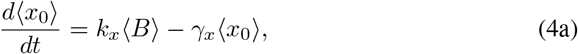

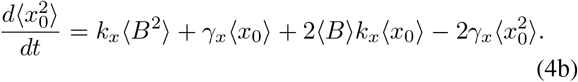

The Fano factor, denoted by *FF* in this paper, is a crucial metric for quantifying random fluctuations in these systems. It is defined as the variance divided by the mean [14], [77], as illustrated below,

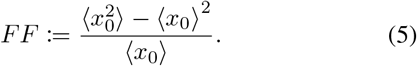

We investigate the no-delay system’s steady-state Fano factor. To do this, we solve the system of equations in Eq. at steady state, which yields the steady-state Fano factor as follows:

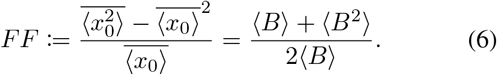

Throughout this article, we focus on the steady-state Fano factor of the active protein level. For a Poisson process, the Fano factor is equal to one because the variance equals the mean. This limit is recovered in our model when protein production is non-bursty, such that the distribution *α*_*j*_ is concentrated at *B* = 1 (i.e., *α*_1_ = 1). To generalize these results, all biochemical rates and time constants are non-dimensionalized by scaling time to the protein half-life. Under this convention, the decay rate is *γ*_*x*_ = 1, which implies that one unit of time represents approximately 1.44 protein half-lives.

## III. One-Step Delay Model

Building on the no-delay model, we introduce a one-step delay system to account for temporal delays in protein activation. This stochastic time-delay system consists of three events:

i. burst production of inactive proteins *X*_0_,
ii. conversion of inactive proteins *X*_0_ to active proteins *X*_1_ through a one-step delay, and
iii. degradation of the active protein *X*_1_.

Inactive proteins *X*_0_ are converted to the active form *X*_1_ via a stochastic process, where the waiting time is exponentially distributed with mean *τ/f* (*x*_1_). The state-dependent delay function, *f*(*x*_1_), modulates the conversion rate based on the current level of active proteins, *x*_1_(*t*). Each event changes the population counts of *x*_0_(*t*) and *x*_1_(*t*), as summarized in Table II. The probability of each event occurring within the infinitesimal time interval (*t, t* + *dt*] is given accordingly. Therefore, the function *f*(*x*_1_) controls the statistical characteristics of the delay process in addition to determining whether the conversion dynamics are accelerated or attenuated.

**TABLE II:** Stochastic gene expression model with one-step state-dependent activation delay.

| Event | Change in Population Count | Probability Event will Occur in $(t, t + dt]$ |
| --- | --- | --- |
| Bursty Protein Synthesis | $x_0 \rightarrow x_0 + j$ | $k_x \alpha_j dt, j \in \{1, 2, \dots\}$ |
| One-Step Delay Conversion | $x_0 \rightarrow x_0 - 1$<br>$x_1 \rightarrow x_1 + 1$ | $\frac{f(x_1)}{\tau} x_0(t) dt$ |
| Protein Degradation | $x_1 \rightarrow x_1 - 1$ | $\gamma_x x_1(t) dt$ |

Using Table II, the differential equations governing the moments of *x*_0_(*t*) and *x*_1_(*t*) are given by:

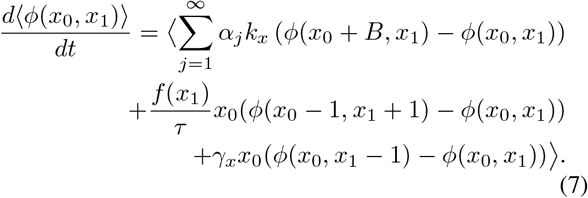

Applying the test function 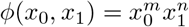 to Eq. (7) with *m* = 1 and *n* = 1 yields the following first-order moments:

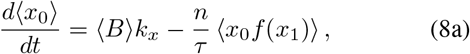

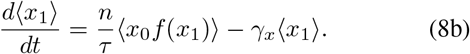

To determine the steady-state mean, we set the right-hand side of the moment equations in Eq. (8) to zero, yielding:

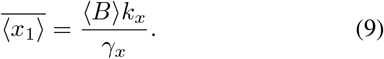

This result shows that the steady-state level of active protein does not depend on the delay or its specific functional form. This occurs because, in the model, the inactive protein cannot be degraded. The part in the propensity function *x*_0_*f*(*x*_1_) in the previous equation introduces nonlinearity into the system due to the multiplicative coupling between the state-dependent delay function *f*(*x*_1_) and the stochastic variable *x*_0_. We linearize it around the steady state 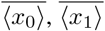 using the LNA [63], [78]–[80], and we get:

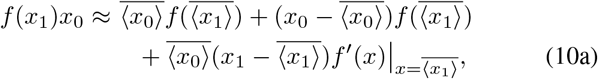

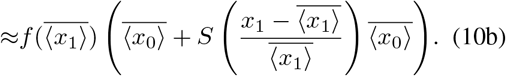

The slope of the function at steady state is represented by 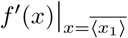. Here, *S* is the logarithmic sensitivity of *f*(*x*_1_) evaluated at the steady-state mean, 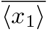, quantifying the relative change of the delay function with respect to variations in *x*_1_ around its steady state,

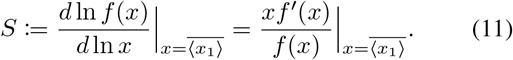

Therefore, we can choose a function of *x*_1_ such that, at the steady state 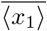, it satisfies the following condition:

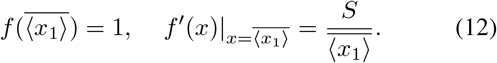

The parameter *S* directly relates this slope to the magnitude and direction of feedback at the mean steady-state level:

- Positive feedback:

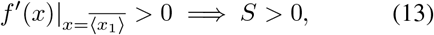

indicating that an increase in the mean number of active proteins accelerates further activation. This can amplify fluctuations or produce switch-like behavior. Biologically, positive feedback represents auto-activation or cooperative signaling mechanisms [81], [82].

- Negative feedback:

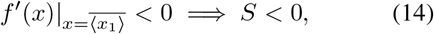

indicating that an increase in the mean number of active proteins slows further activation. Biologically, negative feedback models homeostatic regulation or inhibitory pathways [83], [84].

The first-order steady-state moments are obtained by applying the LNA to Eq. (8) and setting the right-hand side to zero; second-order moments are derived in a similar manner. Those moments are then used for subsequent fluctuation analysis, which gives us the steady-state Fano factor as follows:

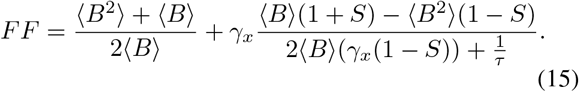

The Fano factor consists of two components: the first term corresponds to the Fano factor of the no-delay system (Eq. (6)), while the second term arises due to the state-dependent activation delay. The steady-state Fano factor depends on *S*, the log-sensitivity of the state-dependent delay function. At steady state, the delay for each activation step is exponentially distributed with mean 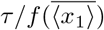. Since 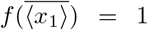 (Eq. (12)), the mean of this exponential distribution reduces to *τ* . The influence of the average activation delay *τ* on steady-state protein fluctuations is illustrated in Fig. 2(B). To ensure bounded fluctuations, the log-sensitivity of the function must remain *S* < 1. Fluctuations rise as *S* increases for a range of delay log-sensitivity values. As the figure shows, fluctuations remain below or equal to those of the no-delay system when the log sensitivity is below the critical value *S*^*c*^. The analytical critical value is,

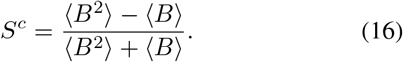

**Fig. 2:**
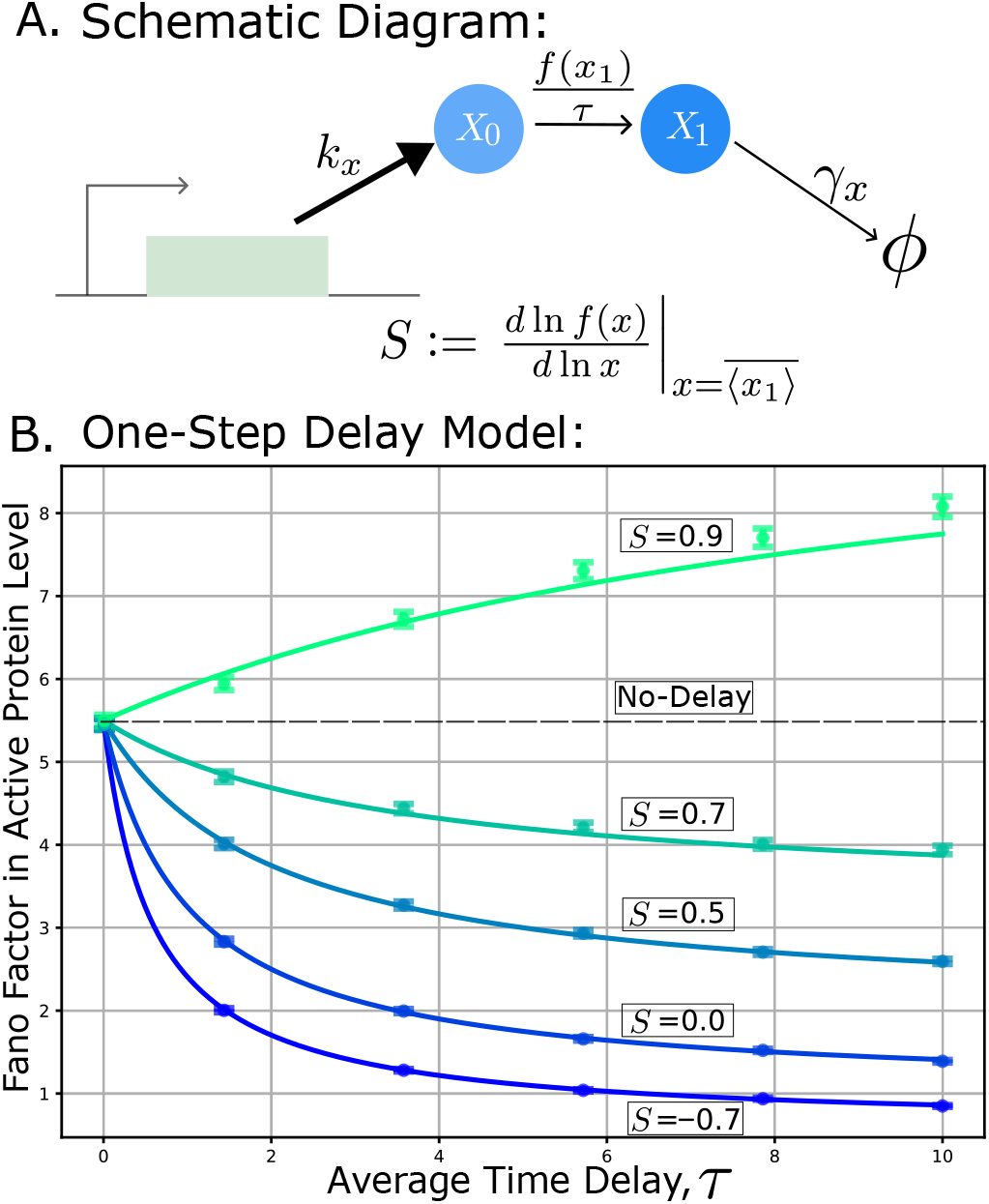
The Fano factor for active protein levels decrease with increasing average activation delay when the log-sensitivity of the state-dependent delay function is below its critical value. **A**. Schematic of the stochastic gene expression model with an one-step state-dependent activation delay, where the delay function *f*(*x*_1_) is determined by the active protein level *x*_1_, and *τ* denotes the average activation delay time (see Table II). **B**. The steady-state active protein fluctuations as a function of the average activation delay time *τ* for the model in panel A: Solid lines represent analytical predictions Eq. (15). Dots indicate the 95% confidence interval using SSA of the gene expression model in Table II. The state-dependent delay function used for the simulation is Eq. (19). The black dotted line denotes the fluctuation level in the no-delay system. For different log sensitivities, *S*, the fluctuations converge to the limiting value given in Eq. 18. The fluctuations can be suppressed below the critical value of S (Eq. (16)). The model parameters are non-dimensionalized by scaling time to the protein half-life, so the decay rate is *γ*_*x*_ = 1. The burst size is *B* = 10, and the burst rate is *k*_*x*_ = 10 per unit time.

For log sensitivities below a critical value, *S*^*c*^, the fluctuations in active protein levels decrease as the average time delay *τ* increases. The system without any delay, implying *τ* = 0, gives us the Fano factor of the no-delay system,

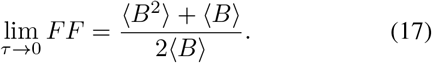

In the limit of an infinitely long delay, the steady-state Fano factor converges to,

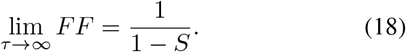

This expression highlights different regimes depending on the value of *S* (Fig. 2(B)):

- For 0 *< S* ≤ *S*^*c*^, the Fano factor decreases as *τ* increases, eventually approaching a value above the Poisson limit (*FF* > 1). In other words, introducing a delay reduces fluctuations compared to the no-delay case, but the fluctuations remain larger than the Poisson Fano factor.
- For *S* ≤ 0, corresponding to negative feedback, the Fano factor can fall below one, indicating that the system can suppress fluctuations below the Poisson limit. For very large negative *S*, the Fano factor can approach zero, meaning fluctuations are strongly suppressed.

From a biological perspective, these findings suggest that cells can regulate stochastic fluctuations in protein levels by exploiting state-dependent delays. The SSA [54] shows the time evolution of protein levels for the no-delay system (light blue solid line) and the one-step delay system (deep blue dashed line) in Fig. 1(B). The corresponding histograms of protein levels reveal reduced fluctuations in the one-step delay system compared to the no-delay system. Negative feedback in the activation delay effectively reduces fluctuations, enabling precise control of protein variability, while moderate positive feedback can also reduce fluctuations compared to a no-delay system. Notably, this regulation can suppress fluctuations below the Poisson limit (*FF* = 1). Overall, the log-sensitivity of the delay emerges as a key mechanism through which cells fine-tune protein-level fluctuations. For SSA, we have used the following functional form:

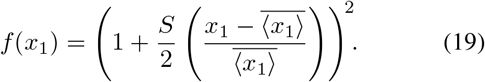

This functional form is chosen to be consistent with the specified sensitivity at steady state. As shown in Figure 2, the SSA of the gene expression model in Table II, incorporating the state-dependent activation delay defined in Eq. (19), closely matches the analytical predictions.

## IV. Multi-Step Delay Model

Many biological processes involve multiple intermediate steps before a protein becomes functionally active, such as folding, modification, or transport. To capture sequential maturation and variability in activation timing, we extend the one-step state-dependent delay model to a multi-step state-dependent delay model, where *n* denotes the number of activation delay stages; small *n* produces a noisy, broadly distributed delay, while large *n* approaches a nearly fixed delay [85]. In the multi-step delay system, the inactive proteins *X*_0_ progress through a series of *n*-delay stages before becoming active proteins *X*_*n*_. Each delay stage represents a first-order stochastic process with a rate 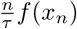 where *f*(*x*_*n*_) is the state-dependent delay function (Fig. 3(A)). Given the current level of active proteins *x*_*n*_, each conversion step is governed by an exponential waiting time. Consequently, the total delay for an inactive protein to progress through all *n* steps follows an Erlang distribution with mean *τ/f* (*x*_*n*_) and coefficient of variation 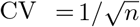. In the limit *n* → ∞, CV → 0 and the delay approaches the deterministic value *τ/f* (*x*_*n*_). Since *f*(*x*_*n*_) depends on the stochastic variable *x*_*n*_, fluctuations in *x*_*n*_ introduce additional variability in the total delay, and the Fano factor depends on the log-sensitivity of *f*(*x*_*n*_). The moment dynamics for the system from Table III gives us,

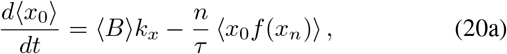

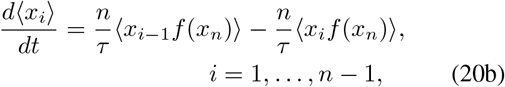

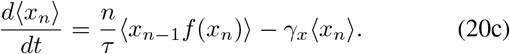

**TABLE III:** Stochastic gene expression model with *n*-step state-dependent activation delay.

| Event | Change in Population Count | Probability Event will Occur in $(t, t + dt]$ |
| --- | --- | --- |
| Bursty Protein Synthesis | $x_0(t) \rightarrow x_0(t) + j$ | $k_x \alpha_j dt, j \in \{1, 2, \dots\}$ |
| One-Step Delay Conversion | $x_0(t) \rightarrow x_0(t) - 1$<br>$x_1(t) \rightarrow x_1(t) + 1$ | $\frac{nf(x_n)}{\tau} x_0(t) dt$ |
| ⋮ | ⋮ | ⋮ |
| $n$-Step Delay Conversion | $x_{n-1} \rightarrow x_{n-1} - 1$<br>$x_n \rightarrow x_n + 1$ | $\frac{nf(x_n)}{\tau} x_{n-1} dt$ |
| Protein Degradation | $x_n(t) \rightarrow x_n(t) - 1$ | $\gamma_x x_n(t) dt$ |

**Fig. 3:**
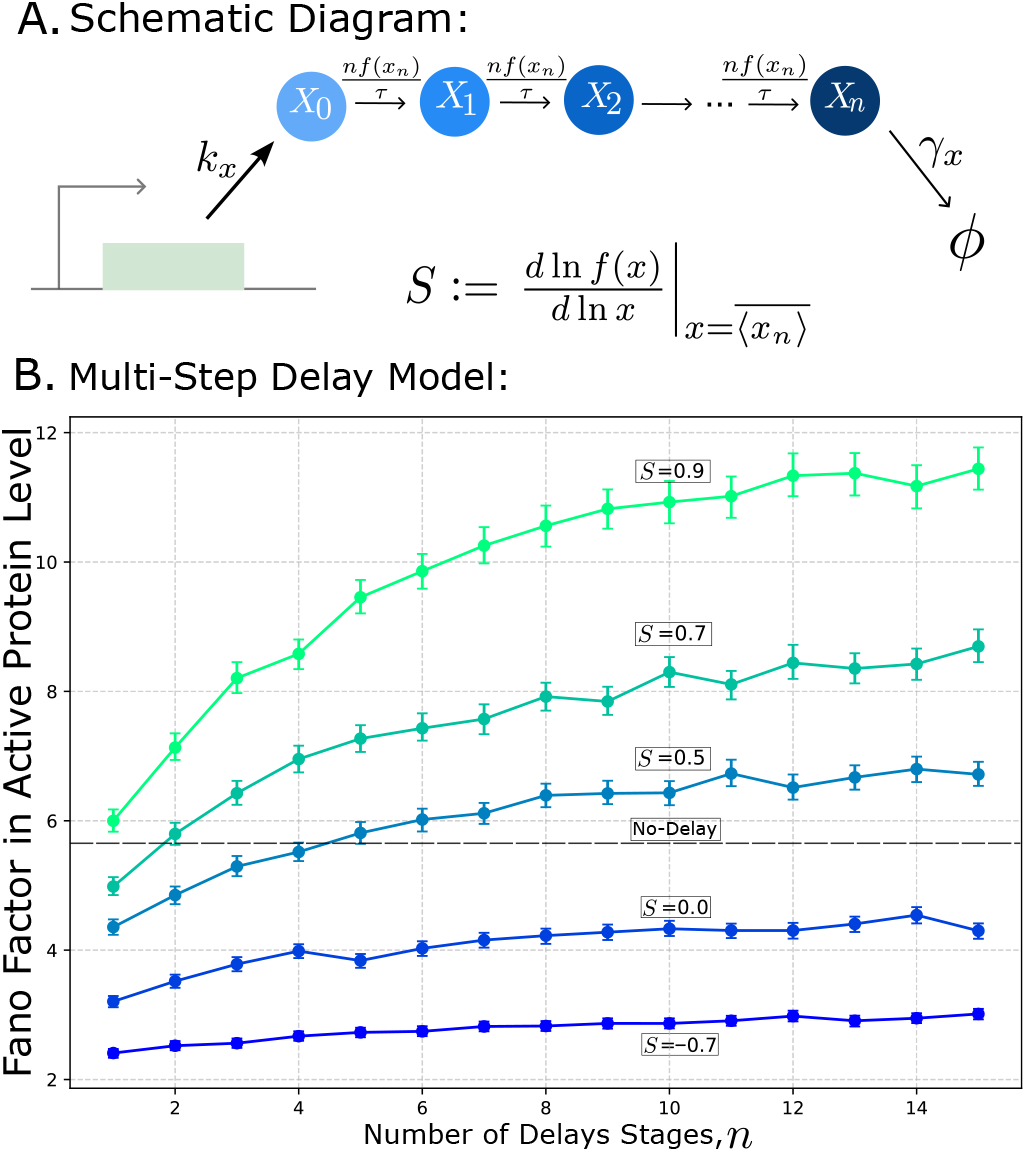
The Fano factor in the active protein level increases with decreasing noise in the state-dependent activation delay. **A**. Schematic of the stochastic gene expression model with an *n*-step state-dependent activation delay, where the delay function *f*(*x*_*n*_) is determined by the active protein level *x*_*n*_, and *τ* denotes the average activation delay time (see Table III). **B**. Fano factor as a function of the number of delay stages, *n* for the model in panel A. Simulations were performed using the SSA of the gene expression model in Table III. The state-dependent function used for the simulation is Eq. (22). The black dashed line corresponds to the Fano factor in the no-delay case. Dots indicate the 95% confidence interval from the simulation. For different log sensitivities, the Fano factor approaches a saturation level after a specific number of delay stages, *n*. Moreover, the Fano factor increases with increasing log sensitivity. The model parameters are non-dimensionalized such that the decay rate is *γ*_*x*_ = 1. Under this scaling, one unit of time corresponds to approximately 1.44 protein half-lives. Accordingly, the burst size is *B* = 10, the burst rate is *k*_*x*_ = 10 per unit time, and an average time delay of *τ* = 1 corresponds to a duration of 1.44 protein half-lives.

For an arbitrary number of delay stages, *n*, the steady-state active protein level can be derived analytically as:

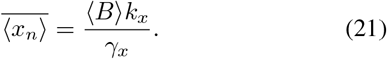

This indicates that active protein levels are independent of both the log-sensitivity parameter *S* and the number of delay stages. The state-dependent activation delay function is chosen as follows, similar to the one-step delay case,

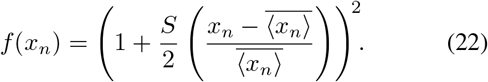

The log-sensitivity expression for the multi-step state-dependent delay model is as follows:

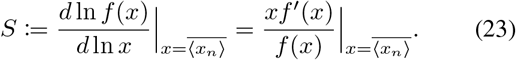

We used the stochastic simulation algorithm to quantify fluctuations in the active protein level. Fixing the total time delay at *τ* = 1, we examined how the number of delay stages, *n*, affects the fluctuations in the active protein level. As shown in Fig. 3(B), an increasing number of delay stages, *n* initially elevates Fano factor, which then saturates after several steps. Notably, for modest values of log-sensitivity (e.g., *S* ≤ 0), the system exhibits a lower Fano factor compared to the no-delay case, suggesting that state-dependent delays can reduce stochastic fluctuations below a certain level of log-sensitivity control even in multi-step delay model.

## V. One-and Multi-Step Delay Models: with Feedback in Burst Frequency

Next, we extend the one-step state-dependent delay model to incorporate negative feedback in burst frequency, a common motif in gene regulatory networks [33], [35], [37],[52], [86], [87]. This trancriptional feedback regulating burst frequency is implemented by having the active protein *X*_1_ inhibiting the protein burst production *X*_0_ as shown in Fig. 4(A). In this model, two different feedback mechanisms are present. One arises from the state-dependent activation delay introduced earlier, and the other is introduced here through the burst frequency. In the stochastic formulation, the burst rate *k*_*x*_ becomes a monotonically decreasing function of the active protein level, *k*_*x*_(*x*_1_(*t*)), implying that higher concentrations of active protein reduce the likelihood of additional burst production. Introducing this feedback renders the system nonlinear and, as a result, precludes an exact analytical solution. We use the feedback function *k*_*x*_(*x*_1_(*t*)) and get the first-order moment of *x*_1_ in the following manner,

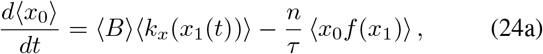

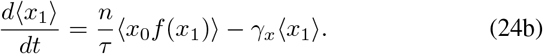

**Fig. 4:**
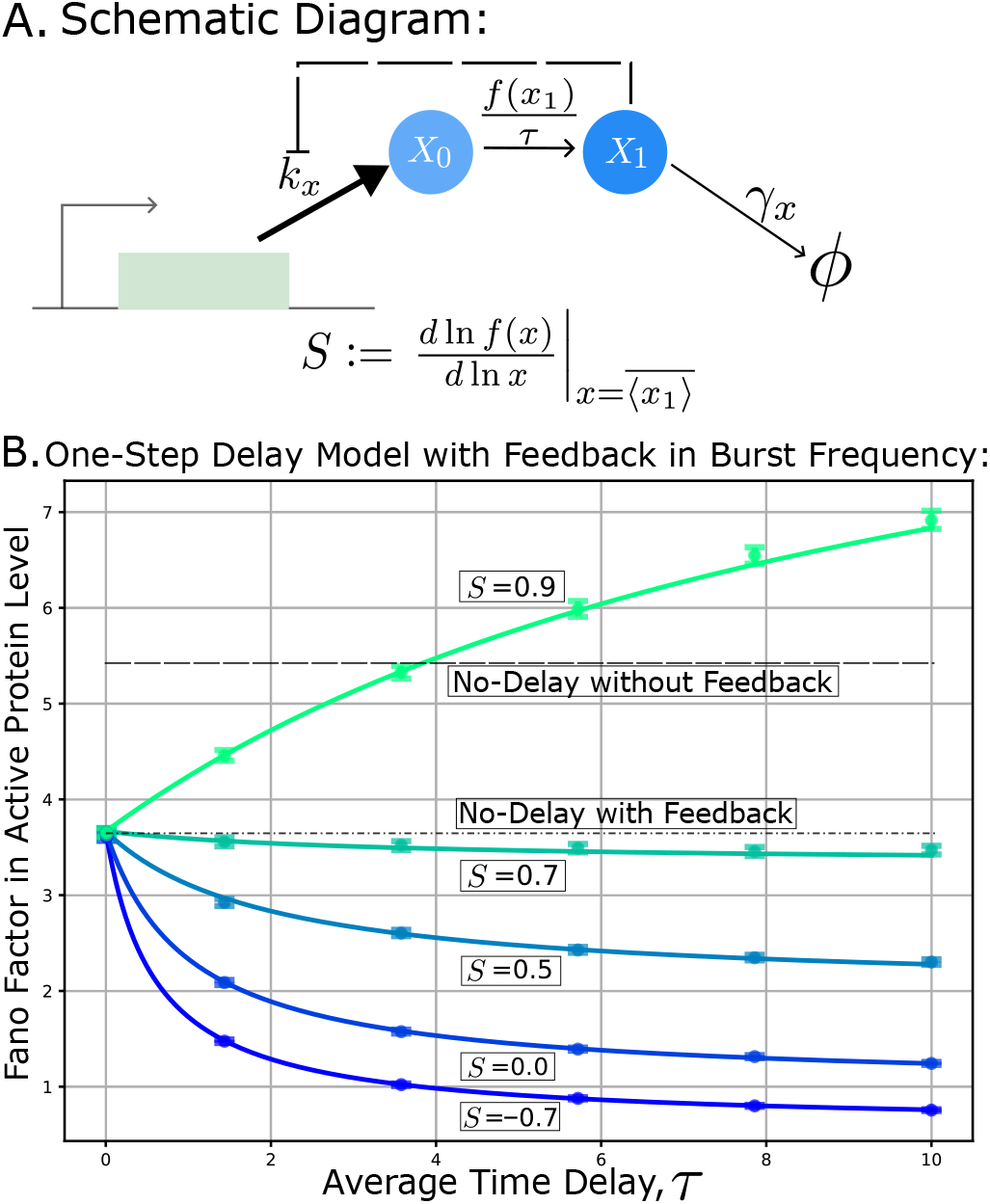
The Fano factor of active protein level decrease as the average activation delay increases, provided the log-sensitivity of the state-dependent delay is below its critical value. **A**. Schematic of the stochastic gene expression model with an one-step state-dependent activation delay, where the delay function *f*(*x*_1_) is determined by the active protein level *x*_1_, and *τ* denotes the average activation delay time (see Table II). The active protein *X*_1_ negatively regulates the burst production rate *k*_*x*_ via a feedback loop (black dotted line). **B**. Steady-state protein fluctuations, *FF* as a function of the average activation delay time *τ* for the model in panel A: The solid lines represent analytical predictions Eq. (28). The dots represent the 95% confidence interval from the SSA of the gene expression model with one-step delay Table II with feedback function Eq.(32). The state-dependent function used for the simulation is Eq. (19). The black dotted line denotes the fluctuation level of the no-delay system without feedback. The dot-dashed line corresponds to the steady-state Fano factor with feedback in burst frequency. For different log sensitivities, *S*, the Fano factor converges to the limiting value given in Eq. (18). The fluctuations can be suppressed below the critical value of *S*^*c*^ (Eq. (29)). The model parameters are: burst size *B* = 10, burst rate *k*_*x*_ = 10 per unit time, decay rate *γ*_*x*_ = 1, and feedback strength *ε* = 0.5.

The steady-state mean for the active protein is,

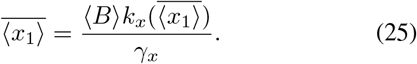

We assume that the above equation admits a unique solution for 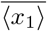. We then consider small fluctuations around the steady-state active protein level and use the LNA to linearize the rate *k*_*x*_(*x*_1_(*t*)) about the steady state as follows:

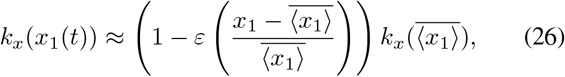

where *ε* is defined as feedback strength [34], [36], a dimensionless constant, and

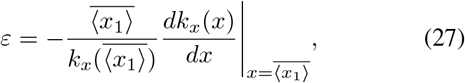

is the log-sensitivity of the burst rate varying depending on the active protein level *x*_1_ evaluated at steady state. Since the LNA assumes small fluctuations around the steady state, we assume that the feedback strength *ε* is sufficiently small so that this approximation remains valid. Following the approach in Section III, we apply the LNA to the state-dependent activation delay. At steady state, this yields the Fano factor for a one-step delay with feedback in burst frequency:

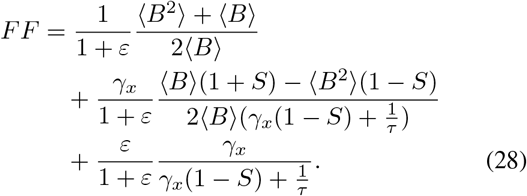

where *S* the log-sensitivity of the state-dependent delay function (Eq. (11)). The Fano factor consists of three components: the first represents the fluctuations arising from feedback in the no-delay system [34], [88], and the rest of the terms arise from the effect of the state-dependent delay with feedback. Increasing the feedback strength *ε* reduces the no-delay term because stronger feedback suppresses fluctuations [34], [36], [87]. In the second term, feedback only appears through the factor 1*/*(1 + *ε*), so its effect is modest for small *ε*. The third term, the coefficient *ε/*(1 + *ε*) is approximately equal to *ε* for small values, so this contribution increases roughly linearly with the feedback strength *ε*. As a result, the overall Fano factor is primarily governed by the reduction in the no-delay term, and the modest increase from the third term slightly offsets this effect, leading to a small net decrease in noise. Similar to Section III, there exists a critical value for the log sensitivity, *S*^*c*^, such that, depending on the feedback strength, the fluctuations remain less than or equal to those in the no-delay system,

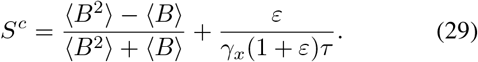

In Eq. (29), the first term corresponds to the critical log sensitivity for the one-step delay system (Eq. (16)), while the second term represents the contribution of negative feedback. For log sensitivity less than *S*^*c*^, the fluctuations decrease as the average time delay, *τ*, increases, as shown in Fig.4(B). The system without any delay, implying *τ* = 0, gives us the Fano factor of the no-delay system with feedback in burst frequency,

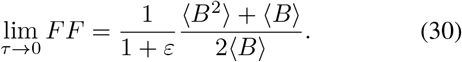

This shows that increasing feedback strength suppresses noise in the active protein level. In the case of infinite time delay, the Fano factor converges to,

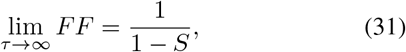

which is equivalent to the Fano factor for the one-step delay model (Eq. (18)). From a biological perspective, this finding suggests that the state-dependent delay and feedback can be adjusted to reduce protein fluctuations. This suggests that cells may use both negative feedback and temporal delays to control fluctuations in protein levels. The feedback function chosen for the simulation is,

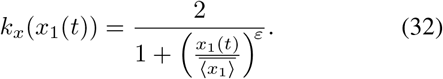

where *ε* is the feedback strength. This form is chosen to be consistent with the specified sensitivity at the steady state. As shown in Figure 4, the SSA of the gene expression model in Table II, incorporating the feedback function defined in Eq. (32), closely matches the analytical predictions.

To investigate how feedback influences fluctuations in systems with *n*-delay stages, we extend the one-step delay model with burst-frequency feedback to a multi-step delay framework. In this case, the active protein *X*_*n*_ prevents the burst production *X*_0_ in this setup shown in Fig. 5(A). The corresponding first-order steady-state moments for the feedback system can be obtained in the same way as for the one-step delay with feedback case. The feedback function chosen for the simulation is,

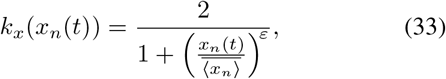

where *ε* is the feedback strength. Feedback in burst frequency does not significantly reduce the Fano factor in the multi-step delay system, as Fig. 5(B) illustrates. The overall Fano factor remains similar to that in the no-feedback scenario. They reach saturation after some specific delay stages. Fluctuations can actually be suppressed below the no-delay scenario for *S* < 0. These findings demonstrate that negative feedback with multiple delay stages suppresses fluctuations more effectively than the no-delay case.

**Fig. 5:**
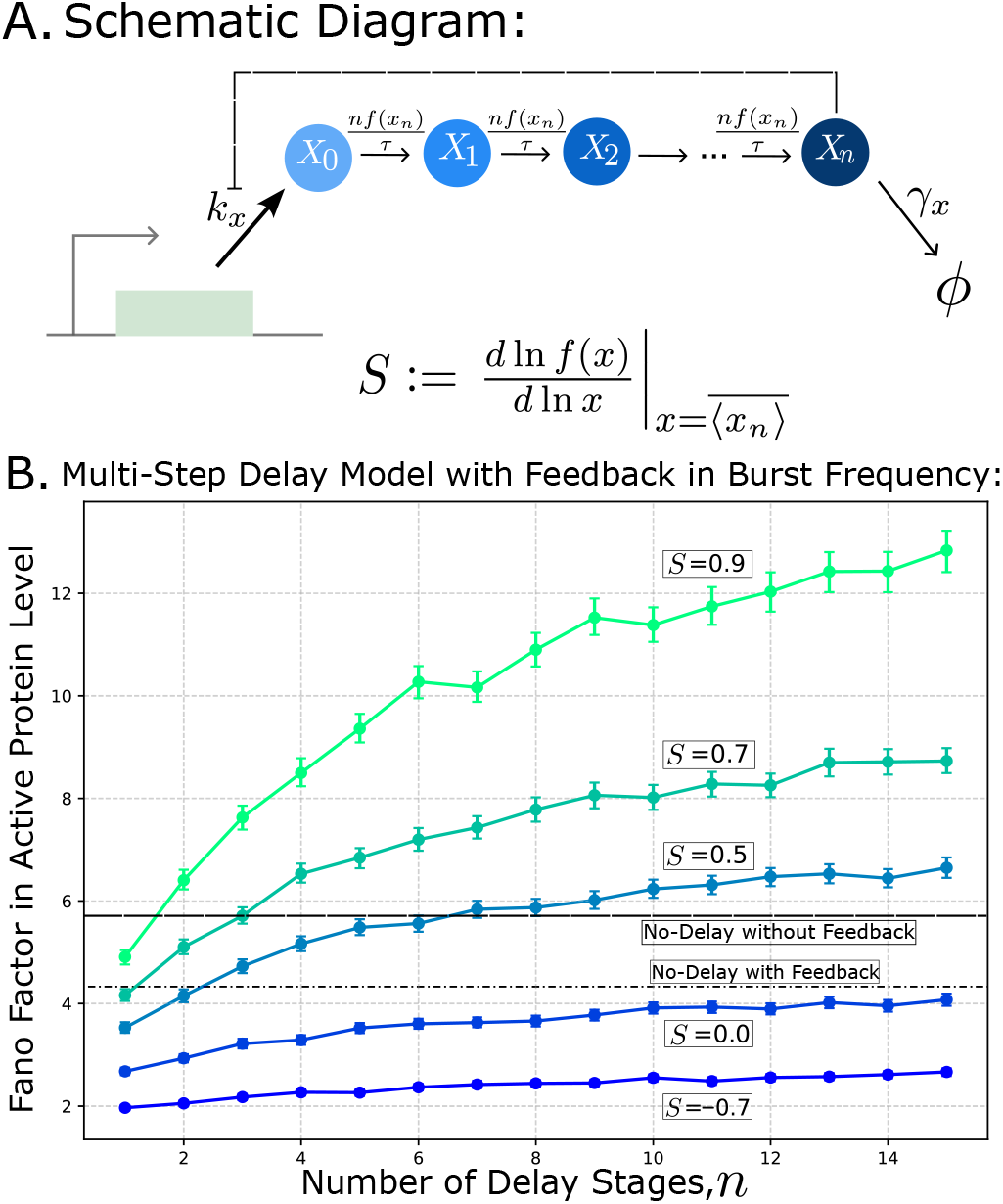
The Fano factor in the active protein level increases with decreasing noise in the state-dependent activation delay. **A**. Schematic of the stochastic gene expression model with an *n*-step state-dependent activation delay, where the delay function *f*(*x*_*n*_) is determined by the active protein level *x*_*n*_, and *τ* denotes the average activation delay time (see Table III). The active protein *X*_*n*_ negatively regulates the burst production rate *k*_*x*_ via a feedback loop (black dotted line). **B**. Fano factor as a function of the number of delay stages, *n*, for the model in panel A with feedback in burst frequency. The black dashed line corresponds to the Fano factor with no-delay without feedback. The dot-dashed line corresponds to the Fano factor of no-delay with feedback in burst frequency. Simulations were performed using the SSA of the gene expression model in Table III with feedback function Eq. (33). The state-dependent function used for the simulation is Eq. (22). The Fano factor reaches a saturation level after a specific number of delay stages, *n*. As the log-sensitivity *S* increases, the Fano factor also increases. However, for *S* ≤ 0, the Fano factor can be suppressed below the no-delay with feedback case. The state-dependent function used for the simulation is Eq. (22). The feedback function is Eq. (33). The model parameters are: burst size *B* = 10, burst rate *k*_*x*_ = 10 per unit time, decay rate *γ*_*x*_ = 1, and feedback strength *ε* = 0.5.

## VI. Discussion

In this study, we examined how stochastic fluctuations in protein expression are shaped by state-dependent delays. We studied a system where *X*_0_ transitions through multiple intermediate stages to become the active protein *X*_*n*_, and the level of *X*_*n*_, which in turn modulates the burst frequency of *X*_0_ production. A fundamental aspect of biology is encapsulated in this formulation: gene expression delays are frequently not fixed but rather rely on the system’s present circumstances [89]. Using both the Linear Noise Approximation (LNA) and the Stochastic Simulation Algorithm (SSA), we were able to quantify the Fano factor in the active protein level across different delay structures and feedback configurations.

Our investigation in Section III reveals that, in contrast to the no-delay system, state-dependent delays do not always increase fluctuations (Fig. 2). Surprisingly, under specific parameter regimes, fluctuations can be reduced below no-delay levels in the presence of negative feedback (Section V). This suggests that state-dependent modulation via delay, in combination with feedback (Fig. 4), can efficiently regulate fluctuations. In biological systems, such behavior might be advantageous for preserving steady protein levels even when there is inherent stochasticity.

We extend the framework to *n*-delay stages and show that the fluctuation properties depend systematically on *n* (Section IV). Biologically, these stages may represent intermediate processes such as transcription, translation, protein folding, or intracellular transport before the final product becomes active. The accumulation of stochastic effects across multiple steps causes the variation in the active protein to increase as the number of delay stages, *n*, increases (Fig. 5). However, the Fano factor converges to a specific saturation level after a certain number of delay stages, *n*. This suggests that in systems with state-dependent delays, Fano factor levels do not diverge indefinitely; instead, they remain bounded even as *n* goes to infinity. This demonstrates that the proposed framework is applicable across a wide range of delay architectures. Overall, these results support the notion that biological systems can regulate stochastic fluctuations while employing multi-step processes such as transcriptional elongation or protein maturation.

Our analysis raises new problems while capturing some important characteristics of delay-dependent regulation. A particular kind of state dependency *f*(*x*_*n*_) with fixed log sensitivity was assumed; however, other nonlinear forms might result in substantially different fluctuations and scaling. Future research should evaluate predictions against single-cell data and expand the current framework to include such complexities. Furthermore, a more thorough knowledge of how delay-dependent feedback functions under actual biological conditions would be possible by taking into account extrinsic noise sources, such as variations in global cellular resources, transcriptional machinery, or environmental variability. Recent single-cell studies in *Saccharomyces cerevisiae* [90] demonstrate how extrinsic factors such as cell size and cellular resources contribute to gene expression variability, providing a framework for incorporating such noise sources into delay-based models. The durability of multi-step delay designs in controlling gene expression variability may be further clarified by considering both intrinsic and extrinsic noise contributions, which may reveal different characteristics.

Overall, our findings provide theoretical evidence that cells can use state-dependent delays to achieve robust fluctuations control. Rather than being detrimental, delays tuned to the system state may serve as an adaptive strategy for balancing responsiveness and stability. This perspective aligns naturally with broader themes in systems biology, where timing and fluctuation regulation play a central role in shaping cellular function.

## VII. Acknowledgments

This work is supported by NIH-NIGMS via grant R35GM148351.

## References

[1] P. S. Swain, M. B. Elowitz, and E. D. Siggia, “Intrinsic and extrinsic contributions to stochasticity in gene expression,” Proceedings of the National Academy of Sciences, vol. 99, no. 20, pp. 12 795–12 800, 2002.

[2] A. Singh and M. Soltani, “Quantifying intrinsic and extrinsic variability in stochastic gene expression models,” PLOS One, vol. 8, no. 12, p. e84301, 2013.

[3] M. Scott, B. Ingalls, and M. Kærn, “Estimations of intrinsic and extrinsic noise in models of nonlinear genetic networks,” Chaos: An Interdisciplinary Journal of Nonlinear Science, vol. 16, no. 2, 2006.

[4] D. Orrell and H. Bolouri, “Control of internal and external noise in genetic regulatory networks,” Journal of Theoretical Biology, vol. 230, no. 3, pp. 301–312, 2004.

[5] J. M. Raser and E. K. O’shea, “Noise in gene expression: origins, consequences, and control,” Science, vol. 309, no. 5743, pp. 2010–2013, 2005.

[6] L. Cai, N. Friedman, and X. S. Xie, “Stochastic protein expression in individual cells at the single molecule level,” Nature, vol. 440, no. 7082, pp. 358–362, 2006.

[7] A. Bar-Even, J. Paulsson, N. Maheshri, M. Carmi, E. O’Shea, Y. Pilpel, and N. Barkai, “Noise in protein expression scales with natural protein abundance,” Nature Genetics, vol. 38, no. 6, pp. 636–643, 2006.

[8] Hilfinger and J. Paulsson, “Separating intrinsic from extrinsic fluctuations in dynamic biological systems,” Proceedings of the National Academy of Sciences, vol. 108, no. 29, pp. 12 167–12 172, 2011.

[9] G. Johnston, B. Gaal, R. P. d. Neves, T. Enver, F. J. Iborra, and N. S. Jones, “Mitochondrial variability as a source of extrinsic cellular noise,” PLOS Computational Biology, vol. 8, no. 3, p. e1002416, 2012.

[10] Zopf, K. Quinn, J. Zeidman, and N. Maheshri, “Cell-cycle dependence of transcription dominates noise in gene expression,” PLOS Computational Biology, vol. 9, no. 7, p. e1003161, 2013.

[11] O. Padovan-Merhar, G. P. Nair, A. G. Biaesch, A. Mayer, S. Scarfone, S. W. Foley, A. R. Wu, L. S. Churchman, A. Singh, and A. Raj, “Single mammalian cells compensate for differences in cellular volume and DNA copy number through independent global transcriptional mechanisms,” Molecular Cell, vol. 58, no. 2, pp. 339–352, 2015.

[12] W. J. Blake, M. Kærn, C. R. Cantor, and J. J. Collins, “Noise in eukaryotic gene expression,” Nature, vol. 422, no. 6932, pp. 633–637, 2003.

[13] Ramsköld G.-J. Hendriks, A. J. Larsson, J. V. Mayr, C. Ziegenhain, M. Hagemann-Jensen, L. Hartmanis, and R. Sandberg, “Single-cell new RNA sequencing reveals principles of transcription at the resolution of individual bursts,” Nature cell biology, vol. 26, no. 10, pp. 1725–1733, 2024.

[14] M. B. Elowitz, A. J. Levine, E. D. Siggia, and P. S. Swain, “Stochastic gene expression in a single cell,” Science, vol. 297, no. 5584, pp. 1183–1186, 2002.

[15] Paulsson, “Models of stochastic gene expression,” Physics of life reviews, vol. 2, no. 2, pp. 157–175, 2005.

[16] Libby, T. J. Perkins, and P. S. Swain, “Noisy information processing through transcriptional regulation,” Proceedings of the National Academy of Sciences, vol. 104, no. 17, pp. 7151–7156, 2007.

[17] Q. Zhang, W. Cao, J. Wang, Y. Yin, R. Sun, Z. Tian, Y. Hu, Y. Tan, and B.-g. Zhang, “Transcriptional bursting dynamics in gene expression,” Frontiers in Genetics, vol. 15, p. 1451461, 2024.

[18] J. Bax, J. Strik, L. A. Schepens, T. L. Lenstra, M. M. Hansen, and H. Marks, “Gene expression noise in development: genome-wide dynamics,” Science & Society, 2025.

[19] Balázsi A. Van Oudenaarden, and J. J. Collins, “Cellular decision making and biological noise: from microbes to mammals,” Cell, vol. 144, no. 6, pp. 910–925, 2011.

[20] A. Raj and A. Van Oudenaarden, “Nature, nurture, or chance: stochastic gene expression and its consequences,” Cell, vol. 135, no. 2, pp. 216–226, 2008.

[21] Singh, B. Razooky, C. D. Cox, M. L. Simpson, and L. S. Weinberger, “Transcriptional bursting from the hiv-1 promoter is a significant source of stochastic noise in hiv-1 gene expression,” Biophysical Journal, vol. 98, no. 8, pp. L32–L34, 2010.

[22] R. D. Dar, S. M. Shaffer, A. Singh, B. S. Razooky, M. L. Simpson, A. Raj, and L. S. Weinberger, “Transcriptional bursting explains the noise–versus–mean relationship in mRNA and protein levels,” PLOS One, vol. 11, no. 7, p. e0158298, 2016.

[23] T. Fukaya, B. Lim, and M. Levine, “Enhancer control of transcriptional bursting,” Cell, vol. 166, no. 2, pp. 358–368, 2016.

[24] J.-W. Veening, W. K. Smits, and O. P. Kuipers, “Bistability, epigenetics, and bet-hedging in bacteria,” Annu. Rev. Microbiol., vol. 62, no. 1, pp. 193–210, 2008.

[25] J. Beaumont, J. Gallie, C. Kost, G. C. Ferguson, and P. B. Rainey, “Experimental evolution of bet hedging,” Nature, vol. 462, no. 7269, pp. 90–93, 2009.

[26] N. Q. Balaban, J. Merrin, R. Chait, L. Kowalik, and S. Leibler, “Bacterial persistence as a phenotypic switch,” Science, vol. 305, no. 5690, pp. 1622–1625, 2004.

[27] S. M. Shaffer, M. C. Dunagin, S. R. Torborg, E. A. Torre, B. Emert, Krepler, M. Beqiri, K. Sproesser, P. A. Brafford, M. Xiao et al., “Rare cell variability and drug-induced reprogramming as a mode of cancer drug resistance,” Nature, vol. 546, no. 7658, pp. 431–435, 2017.

[28] A. Brock, H. Chang, and S. Huang, “Non-genetic heterogeneity—a mutation-independent driving force for the somatic evolution of tumours,” Nature Reviews Genetics, vol. 10, no. 5, pp. 336–342, 2009.

[29] A. A. Chang, J. Jen, S. Jiang, A. Sayad, A. S. Mer, K. R. Brown, M. Nixon, A. Dhabaria, K. H. Tang, D. Venet et al., “Ontogeny and vulnerabilities of drug-tolerant persisters in her2+ breast cancer,” Cancer discovery, vol. 12, no. 4, pp. 1022–1045, 2022.

[30] M. Guha, A. Singh, and N. C. Butzin, “Priestia megaterium cells are primed for surviving lethal doses of antibiotics and chemical stress,” Communications Biology, vol. 8, no. 1, p. 206, 2025.

[31] T. Hossain, A. Singh, and N. C. Butzin, “Escherichia coli cells are primed for survival before lethal antibiotic stress,” Microbiology Spectrum, vol. 11, no. 5, pp. e01 219–23, 2023.

[32] J. Lee, D. H. Swanson, S.-K. Lee, S. Dihardjo, G. Y. Lee, S. Gelle, H. J. Seong, E. R. Bravo, Z. E. Taylor, M. S. Van Nieuwenhze et al., “Trehalose catalytic shift inherently enhances phenotypic heterogeneity and multidrug resistance in mycobacterium tuberculosis,” Nature communications, vol. 16, no. 1, p. 6442, 2025.

[33] A. Singh and J. P. Hespanha, “Evolution of gene auto-regulation in the presence of noise,” IET Systems Biology, vol. 3, no. 5, pp. 368–378, 2009.

[34] Smith and A. Singh, “Noise suppression by stochastic delays in negatively autoregulated gene expression,” in 2020 American Control Conference (ACC). IEEE, 2020, pp. 4270–4275.

[35] Becskei and L. Serrano, “Engineering stability in gene networks by autoregulation,” Nature, vol. 405, no. 6786, pp. 590–593, 2000.

[36] Singh, “Negative feedback through mRNA provides the best control of gene-expression noise,” IEEE Transactions on Nanobioscience, vol. 10, no. 3, pp. 194–200, 2011.

[37] Nevozhay, R. M. Adams, K. F. Murphy, K. Josić, and G. Balázsi, “Negative autoregulation linearizes the dose–response and suppresses the heterogeneity of gene expression,” Proceedings of the National Academy of Sciences, vol. 106, no. 13, pp. 5123–5128, 2009.

[38] P. Czuppon and P. Pfaffelhuber, “Limits of noise for autoregulated gene expression,” Journal of Mathematical Biology, vol. 77, no. 4, pp. 1153–1191, 2018.

[39] Y. Tao, X. Zheng, and Y. Sun, “Effect of feedback regulation on stochastic gene expression,” Journal of Theoretical Biology, vol. 247, no. 4, pp. 827–836, 2007.

[40] Z. Zhang, I. Zabaikina, C. Nieto, Z. Vahdat, P. Bokes, and A. Singh, “Stochastic gene expression in proliferating cells: Differing noise intensity in single-cell and population perspectives,” PLOS Computational Biology, vol. 21, no. 6, p. e1013014, 2025.

[41] C. Jia, L. Y. Wang, G. G. Yin, and M. Q. Zhang, “Single-cell stochastic gene expression kinetics with coupled positive-plus-negative feedback,” Physical Review E, vol. 100, no. 5, p. 052406, 2019.

[42] Lestas, G. Vinnicombe, and J. Paulsson, “Fundamental limits on the suppression of molecular fluctuations,” Nature, vol. 467, no. 7312, pp. 174–178, 2010.

[43] S. Modi, S. Dey, and A. Singh, “Noise suppression in stochastic genetic circuits using PID controllers,” PLOS Computational Biology, vol. 17, no. 7, p. e1009249, 2021.

[44] Filo, S. Kumar, and M. Khammash, “A hierarchy of biomolecular proportional-integral-derivative feedback controllers for robust perfect adaptation and dynamic performance,” Nature Communications, vol. 13, no. 1, p. 2119, 2022.

[45] S. Dey, M. Soltani, and A. Singh, “Enhancement of gene expression noise from transcription factor binding to genomic decoy sites,” Scientific reports, vol. 10, no. 1, p. 9126, 2020.

[46] S. Dey, C. A. Vargas-Garcia, and A. Singh, “Controlling gene-expression variability via sequestration-based feedbacks,” IFAC-PapersOnLine, vol. 58, no. 23, pp. 13–18, 2024.

[47] A. Burger, A. M. Walczak, and P. G. Wolynes, “Influence of decoys on the noise and dynamics of gene expression,” Physical Review E—Statistical, Nonlinear, and Soft Matter Physics, vol. 86, no. 4, p. 041920, 2012.

[48] Singh and P. Bokes, “Consequences of mRNA transport on stochastic variability in protein levels,” Biophysical journal, vol. 103, no. 5, pp. 1087–1096, 2012.

[49] W. Bartelds, Ó. Garćıa-Blay, P. G. Verhagen, E. J. Wubbolts, B. van Sluijs, H. A. Heus, T. F. de Greef, W. T. Huck, and M. M. Hansen, “Noise minimization in cell-free gene expression,” ACS Synthetic Biology, vol. 12, no. 8, pp. 2217–2225, 2023.

[50] Q. Wu and T. Tian, “Stochastic modeling of biochemical systems with multistep reactions using state-dependent time delay,” Scientific Reports, vol. 6, no. 1, p. 31909, 2016.

[51] M. Livingston, J. Kwon, O. Valera, J. A. Saba, N. K. Sinha, Reddy, B. Nelson, C. Wolfe, T. Ha, R. Green et al., “Bursting translation on single mRNAs in live cells,” Molecular Cell, vol. 83, no. 13, pp. 2276–2289, 2023.

[52] Bokes and A. Singh, “Gene expression noise is affected differentially by feedback in burst frequency and burst size,” Journal of mathematical biology, vol. 74, no. 6, pp. 1483–1509, 2017.

[53] Volteras, V. Shahrezaei, and P. Thomas, “Global transcription regulation revealed from dynamical correlations in time-resolved single-cell RNA sequencing,” Cell Systems, vol. 15, no. 8, pp. 694–708, 2024.

[54] T. Gillespie, “A general method for numerically simulating the stochastic time evolution of coupled chemical reactions,” Journal of Computational Physics, vol. 22, no. 4, pp. 403–434, 1976.

[55] D. F. Anderson, “A modified next reaction method for simulating chemical systems with time dependent propensities and delays,” The Journal of Chemical Physics, vol. 127, no. 21, 2007.

[56] Schlicht and G. Winkler, “A delay stochastic process with applications in molecular biology,” Journal of Mathematical Biology, vol. 57, no. 5, pp. 613–648, 2008.

[57] A. S. Ribeiro, “Stochastic and delayed stochastic models of gene expression and regulation,” Mathematical Biosciences, vol. 223, no. 1, pp. 1–11, 2010.

[58] M. Barrio, K. Burrage, A. Leier, and T. Tian, “Oscillatory regulation of Hes1: discrete stochastic delay modelling and simulation,” PLoS Computational Biology, vol. 2, no. 9, p. e117, 2006.

[59] D. Bratsun, D. Volfson, L. S. Tsimring, and J. Hasty, “Delay-induced stochastic oscillations in gene regulation,” Proceedings of the National Academy of Sciences, vol. 102, no. 41, pp. 14 593–14 598, 2005.

[60] P. Bokes, J. R. King, A. T. Wood, and M. Loose, “Exact and approximate distributions of protein and mRNA levels in the low-copy regime of gene expression,” Journal of Mathematical Biology, vol. 64, no. 5, pp. 829–854, 2012.

[61] Gedeon, A. R. Humphries, M. C. Mackey, H.-O. Walther, and Z. Wang, “Operon dynamics with state dependent transcription and/or translation delays,” Journal of Mathematical Biology, vol. 84, no. 1, p. 2, 2022.

[62] A. Gupta and M. Khammash, “Stochastic analysis of feedback control with delays in biochemical networks,” The Journal of the Royal Society Interface, vol. 11, no. 95, p. 20140161, 2014.

[63] Singh and R. Grima, “The linear-noise approximation and moment-closure approximations for stochastic chemical kinetics,” arXiv preprint arXiv:1711.07383, 2017.

[64] N. G. Van Kampen, Stochastic processes in physics and chemistry. Elsevier, 1992, vol. 1.

[65] Elf and M. Ehrenberg, “Fast evaluation of fluctuations in biochemical networks with the linear noise approximation,” Genome Research, vol. 13, no. 11, pp. 2475–2484, 2003.

[66] Kursawe, A. Moneyron, and T. Galla, “Efficient approximations of transcriptional bursting effects on the dynamics of a gene regulatory network,” Journal of the Royal Society Interface, vol. 22, no. 227, 2025.

[67] Golding, J. Paulsson, S. M. Zawilski, and E. C. Cox, “Real-time kinetics of gene activity in individual bacteria,” Cell, vol. 123, no. 6, pp. 1025–1036, 2005.

[68] A. Singh, “Transient changes in intercellular protein variability identify sources of noise in gene expression,” Biophysical Journal, vol. 107, no. 9, pp. 2214–2220, 2014.

[69] N. Friedman, L. Cai, and X. S. Xie, “Linking stochastic dynamics to population distribution: an analytical framework of gene expression,” Physical Review Letters, vol. 97, no. 16, p. 168302, 2006.

[70] V. Shahrezaei and P. S. Swain, “Analytical distributions for stochastic gene expression,” Proceedings of the National Academy of Sciences, vol. 105, no. 45, pp. 17 256–17 261, 2008.

[71] Gupta and J. J. Cai, “Gene function revealed at the moment of stochastic gene silencing,” Communications Biology, vol. 8, no. 1, p. 88, 2025.

[72] A. Singh and J. P. Hespanha, “Approximate moment dynamics for chemically reacting systems,” IEEE Transactions on Automatic Control, vol. 56, no. 2, pp. 414–418, 2010.

[73] P. Hespanha and A. Singh, “Stochastic models for chemically reacting systems using polynomial stochastic hybrid systems,” International Journal of Robust and Nonlinear Control: IFAC-Affiliated Journal, vol. 15, no. 15, pp. 669–689, 2005.

[74] Sinigh and J. P. Hespanha, “Stochastic analysis of gene regulatory networks using moment closure,” in 2007 American Control Conference. IEEE, 2007, pp. 1299–1304.

[75] B. Dynkin, “Markov processes,” in Markov Processes: Volume 1. Springer, 1965, pp. 77–104.

[76] B. Dynkin, “Markov processes and semigroups of operators,” Theory of Probability & Its Applications, vol. 1, no. 1, pp. 22–33, 1956.

[77] Paulsson, “Summing up the noise in gene networks,” Nature, vol. 427, no. 6973, pp. 415–418, 2004.

[78] N. G. Van Kampen and W. P. Reinhardt, “Stochastic processes in physics and chemistry,” 1983.

[79] Munsky, W. S. Hlavacek, and L. S. Tsimring, Quantitative Biology: Theory, Computational Methods, and Models. MIT Press, 2018.

[80] S. Modi, M. Soltani, and A. Singh, “Linear noise approximation for a class of piecewise deterministic markov processes,” in 2018 Annual American Control Conference (ACC). IEEE, 2018, pp. 1993–1998.

[81] S.-Y. Shin, O. Rath, S.-M. Choo, F. Fee, B. McFerran, W. Kolch, and K.-H. Cho, “Positive-and negative-feedback regulations coordinate the dynamic behavior of the ras-raf-mek-erk signal transduction pathway,” Journal of Cell Science, vol. 122, no. 3, pp. 425–435, 2009.

[82] O. Cinquin and J. Demongeot, “Roles of positive and negative feedback in biological systems,” Comptes Rendus. Biologies, vol. 325, no. 11, pp. 1085–1095, 2002.

[83] Turrigiano, “Homeostatic signaling: the positive side of negative feedback,” Current Opinion in Neurobiology, vol. 17, no. 3, pp. 318–324, 2007.

[84] Lake, S. A. Corrêa, and J. Müller, “Negative feedback regulation of the erk1/2 mapk pathway,” Cellular and Molecular Life Sciences, vol. 73, no. 23, pp. 4397–4413, 2016.

[85] E. Weidemann, J. Holehouse, A. Singh, R. Grima, and S. Hauf, “The minimal intrinsic stochasticity of constitutively expressed eukaryotic genes is sub-poissonian,” Science Advances, vol. 9, no. 32, p. eadh5138, 2023.

[86] Alon, “Network motifs: theory and experimental approaches,” Nature Reviews Genetics, vol. 8, no. 6, pp. 450–461, 2007.

[87] A. Singh and J. P. Hespanha, “Optimal feedback strength for noise suppression in autoregulatory gene networks,” Biophysical Journal, vol. 96, no. 10, pp. 4013–4023, 2009.

[88] Smith, M. Soltani, R. Kulkarni, and A. Singh, “Modulation of stochastic gene expression by nuclear export processes,” in 2021 60th IEEE Conference on Decision and Control (CDC). IEEE, 2021, pp. 655–660.

[89] Ahmed and E. I. Verriest, “Modeling & analysis of gene expression as a nonlinear feedback problem with state-dependent delay,” IFAC-PapersOnLine, vol. 50, no. 1, pp. 12 679–12 684, 2017.

[90] S. Das, A. Singh, and P. Shah, “Evaluating single-cell variability in proteasomal decay within Saccharomyces cerevisiae,” Biophysical Journal, vol. 124, no. 23, pp. 4072–4086, 2025.

